# Structural role for DNA Ligase IV in promoting the fidelity of non-homologous end joining

**DOI:** 10.1101/2022.10.26.513880

**Authors:** Benjamin M. Stinson, Sean M. Carney, Johannes C. Walter, Joseph J. Loparo

**Author notes:** Correspondence (J.J.L.).

## Abstract

Nonhomologous end joining (NHEJ) is the primary pathway of vertebrate DNA double-strand-break repair. NHEJ polymerases and nucleases can modify DNA ends to render them compatible for ligation, but these enzymes are usually deployed only when necessary for repair of damaged DNA ends, thereby minimizing mutagenesis. Using frog egg extracts, we reveal a structural role for the NHEJ-specific DNA Ligase IV (Lig4) in promoting NHEJ fidelity. Mutational analysis demonstrates that Lig4 must bind DNA ends to form the short-range synaptic complex, in which DNA ends are closely aligned prior to ligation. Furthermore, single-molecule experiments show that a single Lig4 binds both DNA ends at the instant of short-range synapsis. In this way, compatible ends can be rapidly ligated without polymerase or nuclease activity, which we previously showed is restricted to the short-range synaptic complex. Our results provide a molecular basis for the fidelity of NHEJ.

## INTRODUCTION

Faithful repair of DNA double strand breaks (DSBs) is critical for genome stability and tumor suppression. Cells employ two major pathways to repair DSBs: homologous recombination (HR) and non-homologous end joining (NHEJ) (Scully et al. 2019). HR uses a homologous template to ensure accurate restoration of the DNA sequence. In contrast, NHEJ directly re-ligates broken DNA ends without a template and thus is able to function in all phases of the cell cycle. Broken DNA ends are often damaged, which renders them incompatible for re-ligation. To overcome this problem, NHEJ utilizes an array of end processing enzymes, including polymerases and nucleases, to resolve incompatible ends and allow ligation (Stinson & Loparo 2021). Such end processing is potentially mutagenic, but accumulating evidence suggests that it is restricted to incompatible ends, and undamaged compatible ends (i.e., blunt or “sticky”) are typically joined without processing (Baumann & West 1998; Labhart 1999; Feldmann et al. 2000; Lin et al. 2013; Waters et al. 2014; Stinson et al. 2020). Thus, ligation is prioritized over end processing, thereby minimizing errors. Here, we reveal the molecular mechanisms underpinning this conservative feature of NHEJ by interrogating how the NHEJ-specific DNA Ligase IV (Lig4) interacts with DNA ends in real time during DSB repair.

The ring-shaped Ku70/80 heterodimer (Ku) initiates NHEJ by encircling broken DNA ends (Walker et al. 2001). Ku serves as a recruitment hub for other NHEJ factors, including the DNA-dependent protein kinase catalytic subunit (DNA-PKcs), XRCC4-like factor (XLF), paralog of XLF and XRCC4 (PAXX), and a complex of XRCC4 and DNA Ligase IV (Lig4), which ultimately joins the DNA ends (Stinson & Loparo 2021). Recent single-molecule and structural studies have elucidated how these core NHEJ factors align DNA ends in preparation for ligation, a process termed synapsis (Graham et al. 2016; Wang et al. 2018; Chen, Lee, et al. 2021). DNA ends are initially held in a “long-range” synaptic complex (LR complex), in which the DNA ends are tethered but not directly juxtaposed (Graham et al. 2016). Ku and DNA-PKcs are sufficient for LR complex formation, but other LR complexes additionally incorporating PAXX and/or XLF have been observed (Wang et al. 2018; Chaplin, Hardwick, Stavridi, et al. 2021). The LR-complex transitions into a “short-range” synaptic complex (SR complex), in which the DNA ends are closely aligned for ligation. Efficient SR-complex formation requires DNA-PKcs catalytic activity, a single XLF homodimer, and the Lig4-XRCC4 complex (Graham et al. 2016). Notably, catalytically inactive Lig4 supports SR-complex assembly (Cottarel et al. 2013; Graham et al. 2016) and even promotes subsequent DSB repair by other factors (Goff et al. 2022), suggesting that Lig4 plays a critical yet ill-defined structural role in SR complex formation.

We previously described how DNA end processing is coordinated with synapsis (Stinson et al. 2020). Using *Xenopus laevis* egg extracts, which efficiently recapitulate many properties of NHEJ observed in cells, we demonstrated that end processing by NHEJ polymerases, nucleases, and other enzymes is largely restricted to the SR-complex. From this observation, we proposed that because Lig4 is required to form the SR-complex, compatible ends are ligated before processing occurs. A critical but untested assumption of this model is that Lig4 binds DNA ends in a ligation-competent state during initial SR complex formation.

Here, we investigate the structural role of Lig4 during SR-complex formation. Using *Xenopus laevis* egg extracts, we show that Lig4 directly binds both DNA ends to assemble the SR-complex. Mutational analysis demonstrates that SR-complex formation requires Lig4 DNA binding. Additionally, single-molecule Förster resonance energy transfer (smFRET) experiments that simultaneously monitor DNA end synapsis and Lig4-DNA binding demonstrate that Lig4 transiently binds DNA ends prior to SR synapsis, and a single Lig4 binds both DNA ends at the onset of SR synapsis. Thus, Lig4 is poised to ligate compatible ends immediately upon formation of the short-range synaptic complex, thereby minimizing errors arising from unnecessary end processing in the SR complex.

## RESULTS

### Characterization of Lig4 DNA-binding mutants

To test whether the Lig4-DNA interaction contributes to synapsis, we generated mutations in the DNA-binding domain (DBD) of Lig4. We selected six basic residues in the DBD of the *X. laevis* Lig4 ortholog that are expected, based on human Lig4-DNA structures (Kaminski et al. 2018), to make electrostatic interactions with DNA: K33, K35, K37, K167, R168, and K169 (Figure 1A). We expressed and purified Lig4-XRCC4 (Lig4-X4) complexes with multiple charge swap mutations: Lig4^mDBD1^ (K33E, K35E, K37E), Lig4^mDBD2^ (K167E, R168E, and K169E) and Lig4^mDBD1+2^ (all six basic→Glu mutations) (Figure S1A). In addition, we purified a Lig4-X4 complex lacking the DBD (Lig4^ΔDBD^; Figure S1A). We assessed DNA binding by these variants using a filter binding assay. Varying concentrations of Lig4-X4 variants were incubated with a ^32^P-labeled ∼1kb circular DNA, and samples were passed sequentially through nitrocellulose and positively charged nylon membranes. In this way, Lig4-X4-DNA complexes were first captured on the nitrocellulose membrane, and free DNA was captured on the nylon membrane to allow calculation of fractional DNA binding (Wong & Lohman 1993). Lig4^mDBD1^ and Lig4^mDBD2^ bound DNA ∼10-fold less strongly than wild-type Lig4, and Lig4^mDBD1+2^ and Lig4^ΔDBD^ bound DNA ∼30-fold less strongly than wild-type (Figure 1B, S1B). Residual DNA binding even in the absence of the Lig4 DBD was likely mediated by the Lig4 catalytic and oligonucleotide/oligosaccharide-fold (OBD) domains, both of which interact directly with DNA (Kaminski et al. 2018). All Lig4 variants showed similar auto-adenylation activity (Figure S1C), suggesting that the mutations introduced do not globally disrupt protein folding. We then tested the ability of the Lig4 variants to support NHEJ. We immunodepleted egg extracts of endogenous Lig4-X4 using an anti-XRCC4 antibody, which co-depletes Lig4 (Graham et al. 2016). Immunodepleted extracts were supplemented with recombinant Lig4-X4 variants. Addition of radiolabeled linear, blunt-ended DNA followed by agarose electrophoresis and autoradiography allowed visualization of end joining. NHEJ kinetics supported by the Lig4-X4 variants mirrored DNA binding affinity: Lig4^mDBD1^ and Lig4^mDBD2^ were modestly defective in end joining, and Lig4^mDBD1+2^ and Lig4^ΔDBD^ were severely defective (Figure 1C). Together, these results establish that Lig4^mDBD1^, Lig4^mDBD2^, and Lig4^mDBD1+2^ are deficient in DNA binding and end joining.

**Figure 1:**
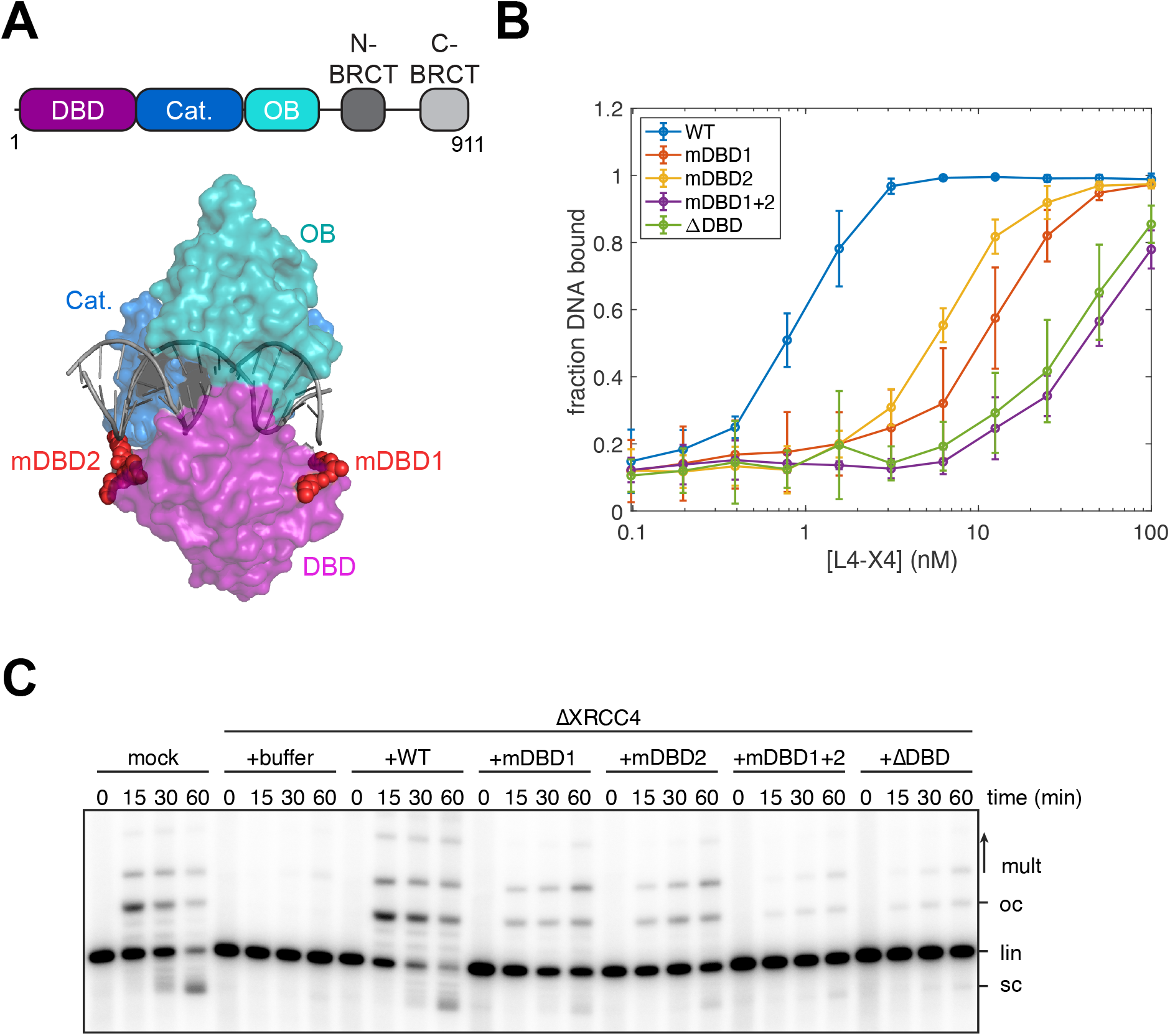
Characterization of Lig4 DNA binding mutants. (A) Domain structure of Xenopus Lig4 and X-ray crystal structure (PDB: 6BKG) of human Lig4. DBD: DNA-binding domain; Cat.: catalytic (adenylation) domain; OB: oligonucleotide/oligosaccharide-binding fold domain. DBD point mutants identified in this study are highlighted in red. (B) Filter binding assay for DNA binding by Lig4 mutants, as detailed in Methods and Figure S1B. Error bars represent standard deviation from three independent experiments. (C) Radiolabeled, blunt-ended, linear DNA molecules were added to the indicated extracts, and reaction samples were stopped at the indicated timepoints. Samples were analyzed by agarose gel electrophoresis and autoradiography. lin: linear; sc: supercoiled; oc: open circular; mult: multimers. Three independent experiments were performed, and a representative autoradiogram is shown.

### DNA-binding by Lig4 is required for short-range synapsis

We next asked whether the Lig4-DNA interaction is required for SR complex assembly. To this end, we tested the Lig4 DNA-binding mutants in single-molecule Förster resonance energy transfer (smFRET) experiments that directly measure short-range synaptic complex formation (Graham et al. 2016). We labeled a ∼3kb blunt-ended DNA fragment with a Cy3B donor fluorophore near one end and a Cy5 acceptor fluorophore near the other (Figure 2A). This DNA substrate was immobilized in a microfluidic flow cell on a streptavidin-functionalized coverslip via an internal biotin linkage. NHEJ was initiated by injecting egg extracts into the flow cell, and juxtaposition of DNA ends in the short-range synaptic complex was monitored by FRET (Figure 2B). As reported previously (Graham et al. 2016), extracts depleted of Lig4-X4 showed severely inhibited short-range synapsis relative to mock-depleted extracts (Figure 2C). Whereas addition of recombinant wild-type Lig4 fully rescued short-range synapsis, addition of each DNA-binding defective mutant showed strongly suppressed short-range synapsis (Figure 2C). Lig4^mDBD1^ and Lig4^mDBD2^ appeared to rescue synapsis to a slightly greater extent than Lig4^mDBD1+2^, although this difference did not rise to the level of statistical significance (Figure 2C). These results demonstrate that the Lig4-DNA interaction is critical for initial formation of the short-range synaptic complex.

**Figure 2:**
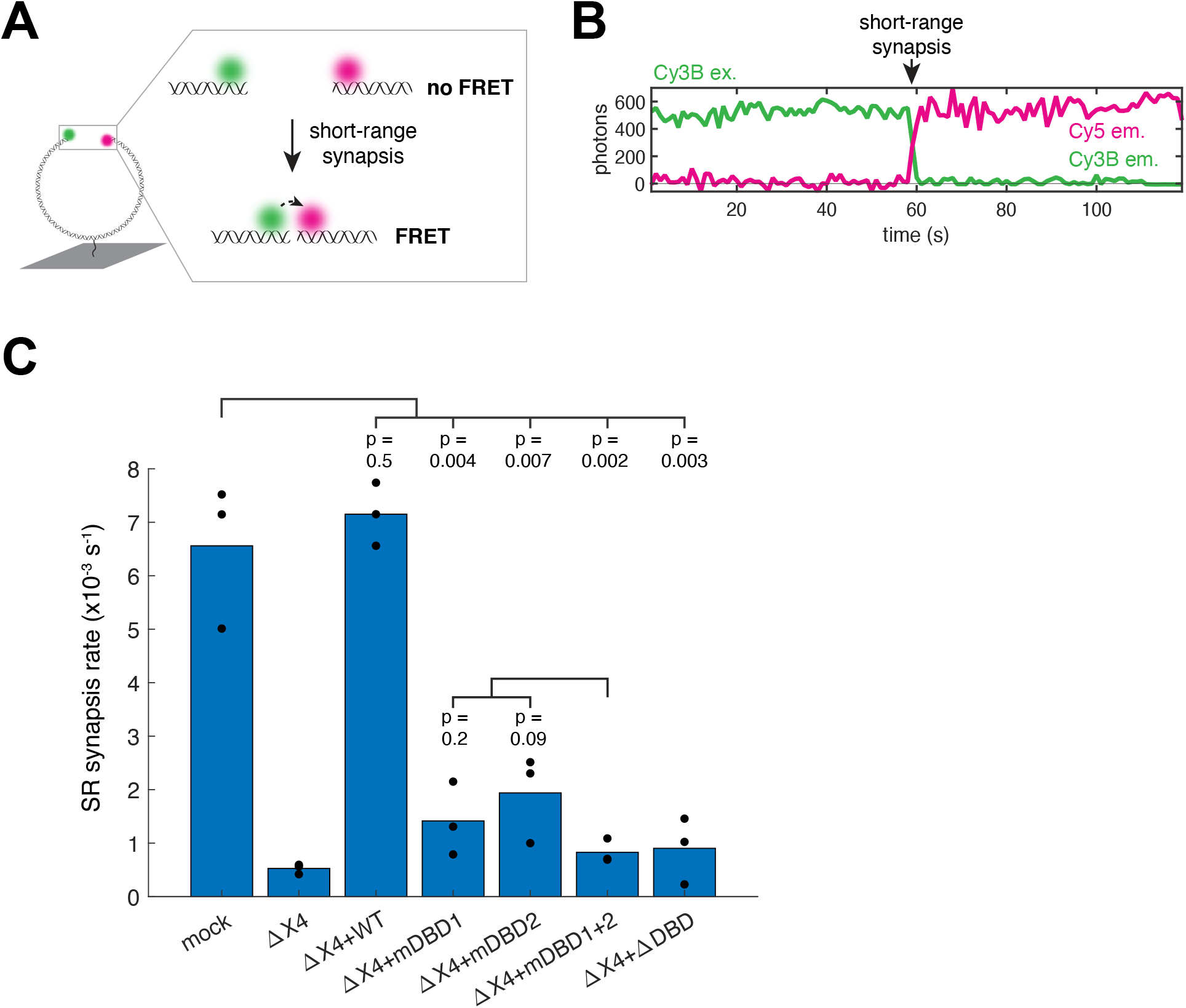
DNA-binding by Lig4 is required for short-range synapsis. (A) Cartoon of smFRET assay for short-range synapsis. Green circle: Cy3B fluorphore; magenta circle: Cy5 fluorophore; dotted arrow: energy transfer. (B) Representative single-molecule trajectory under Cy3B excitation depicting shortrange synapsis. Green line: Cy3B emission; magenta line: Cy5 emission. (C) Extracts were immunodepleted of Lig4-X4 and supplemented with indicated recombinant Lig4-X4 variants, and the rate of short-range synapsis was measured using the assay depicted in panel (A). Black dots: rates from three independent experiments; blue bars: average rates. *p*-values were calculated using the twosample *t*-test.

### DNA binding is required for Lig4 colocalization

To probe the dynamics of Lig4 recruitment and end binding during repair, we designed an smFRET assay to monitor the Lig4-DNA interaction directly (Figure 3A). We modified the DNA substrate shown in Figure 2A to include a Cy3B fluorophore at each DNA end, and we attached a Cy5 fluorophore to an N-terminal yBBR tag on Lig4 (Figure 3A) (Yin et al. 2006). Based on atomic-resolution structures of Lig4 bound to DNA (Kaminski et al. 2018; Chen, Lee, et al. 2021), we expected that end binding by Lig4 would result in FRET from Cy3B to Cy5, whereas recruitment to the overall NHEJ complex without direct DNA binding would result in Cy5 colocalization without FRET (Figure 3A). Extracts were depleted of endogenous Lig4-X4, supplemented with wild-type introduced to the flow cell containing immobilized Cy3B-labeled DNA. An example single-molecule trajectory is shown in Figure 3B, in which the top panel shows Lig4 colocalization with the substrate (Cy5 ex./Cy5 em.), and the middle and bottom panels show Lig4-DNA binding (Cy3B ex./Cy3B and Cy5 em. (middle); calculated FRET_E_ (bottom)). Wild-type Cy5-labeled Lig4-X4, which supported efficient end joining (Figure S2A), showed robust colocalization to DNA spots and minimal binding to coverslip locations that lacked a DNA signal, indicating that the observed colocalization events were highly specific (Figure S2B). ∼70% of colocalization events exhibited Cy3B/Cy5 FRET (FRET_E_ > 0.1), suggesting that wild-type Lig4 directly binds DNA ends for most colocalization events (Figure 3C, WT). To verify that FRET resulted from Lig4-DNA binding, we repeated this experiment with Lig4 DNA-binding mutants. In all cases, perturbing the Lig4-DNA interaction resulted in a lower proportion of high-FRET colocalization events, as well as lower overall colocalization frequency (Figure 3C). Colocalization frequency and the proportion of high-FRET events followed a similar trend as DNA binding, end joining, and rate of short-range synapsis (Figures 1B, 1C, and 2C; WT > mDBD1 ≈ mBDB2 > mDBD1+2 ≈ ΔDBD). These results validate Cy3B/Cy5 FRET in this assay as a reporter of Lig4-DNA binding, and they provide direct evidence that DNA binding is critical for Lig4-X4 colocalization to the NHEJ complex.

**Figure 3:**
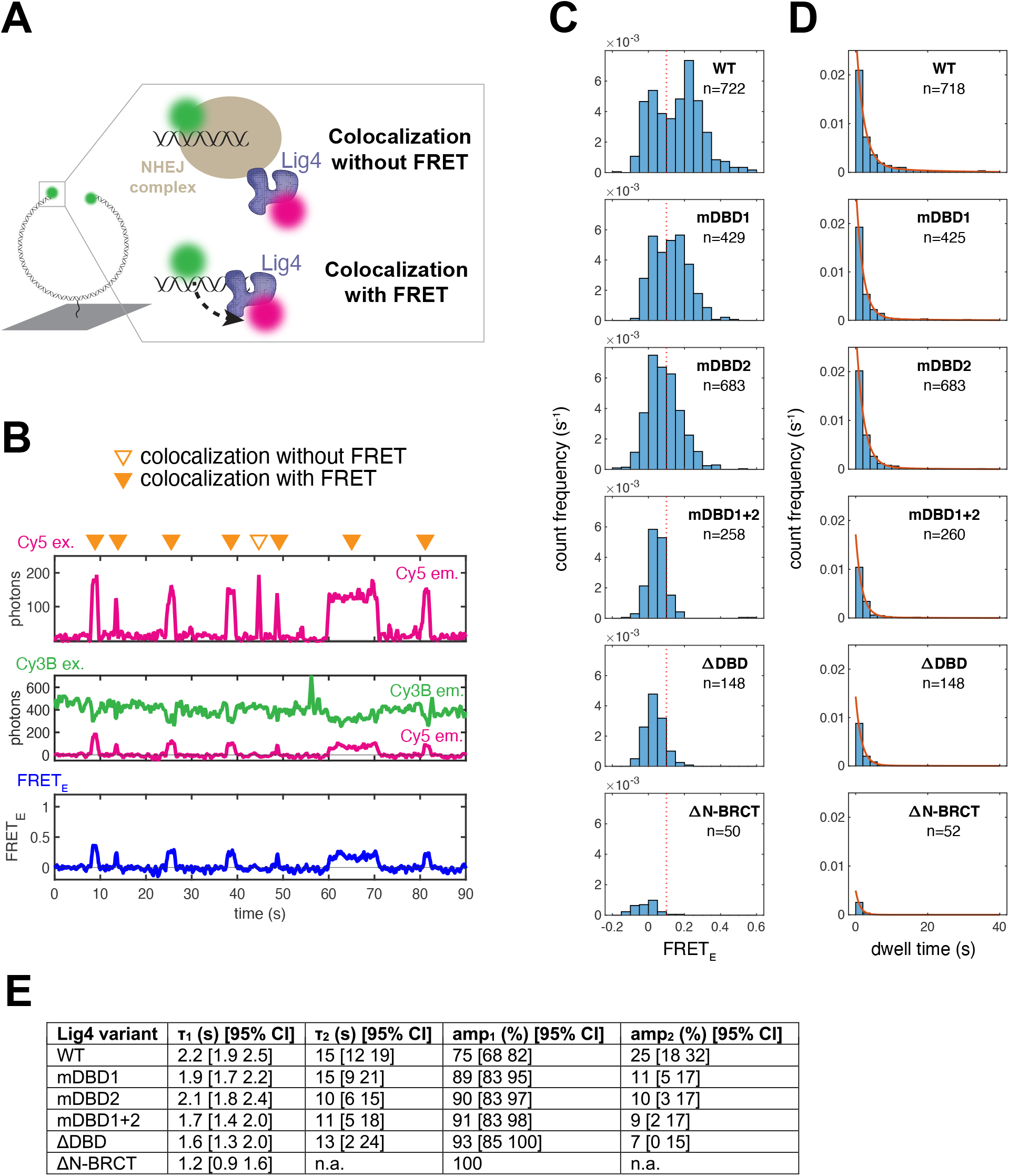
DNA binding is required for Lig4 colocalization. (A) Cartoon of smFRET assay for DNA end binding by Lig4. Green circles: Cy3B fluorophore; magenta circles: Cy5 fluorophore; dotted arrow: energy transfer. Colocalization occurs whenever Lig4 is recruited to the NHEJ complex, but FRET occurs only when Lig4 directly binds DNA ends. (B) Representative single-molecule trajectories showing Lig4 colocalization and end binding. Top panel: Cy5 excitation with Cy5 emission in magenta, measuring Lig4 colocalization; middle panel: Cy3B excitation with Cy3B emission in green and Cy5 emission in magenta, measuring Lig4 DNA binding; bottom panel: calculated FRET efficiency from middle panel. Orange arrows: colocalization events with or without FRET. (C) Colocalization events for each Lig4 variant were detected as described in Methods, and the average FRET_E_ for each colocalization event was calculated and plotted on histograms for each Lig4 variant. Bin counts were normalized by total observation time, such that the y-axis reflects frequency of colocalization events. Data from three independent experiments for each Lig4 variant. (D) Histograms showing dwell time distributions for Lig4 variant colocalization events, detected as described in Methods. Bin counts were normalized by total observation time, such that the y-axis reflects frequency of colocalization events. Red lines show dwell time distribution fits generated by maximum likelihood estimation in MATLAB using one- or two-term exponential models, with fit parameters noted in (E). Data from three independent experiments for each Lig4 variant. (E) Fit parameters for dwell time distributions in (C). Figures in brackets indicate 95% confidence intervals. All distributions were fit to two-term exponential models, except ΔN-BRCT dwells, which were fit to a one-term exponential model. τ, exponential time constant; amp., amplitude.

To address whether Lig4 binds DNA ends before or after short-range synapsis, we treated extracts with the DNA-PKcs inhibitor NU7441, which blocks formation of the short-range synaptic complex (Graham et al. 2016). Strikingly, Lig4-X4 colocalization and end binding was essentially unaltered relative to the DMSO control (Figure S2C). Therefore, the vast majority of Lig4-DNA binding events observed occurred independently of short-range synapsis. Taken together, our results suggest that Lig4 frequently binds DNA ends prior to short-range synapsis.

We next assessed the role of DNA binding in Lig4 colocalization dwell times. For wild-type Lig4, dwell times were biphasic, with a fast phase (time constant ∼ 2 s) accounting for ∼75% of the distribution and a slower phase (time constant ∼15 s) accounting for the remainder (Figure 3D-E; Figure S2D shows single-phase fit). The presence of more than one phase likely reflects distinct populations of Lig4-X4 that make different interactions with other NHEJ factors (see Discussion). Dwell times were similar when Cy5 illumination power was increased (Figure S2E), suggesting that photobleaching did not substantially contribute to the observed kinetics. Disruption of Lig4-DNA binding resulted in less frequent Lig4 recruitment with fewer long dwell times (Figure 3D, amp_2_ in Figure 3E). These results suggest that DNA binding is critical for stable Lig4-X4 retention in the NHEJ complex. Consistent with this idea, wild-type Lig4 colocalization events with high FRET_E_ (>0.1) were longer-lived than low FRET_E_ colocalization events (Figure S2F). Overall, these results show that, despite the presence of multiple protein-protein interactions that are thought to recruit Lig4-X4 to the NHEJ complex (see Discussion), Lig4 colocalization with the NHEJ complex is transient and dependent on DNA binding.

A region in the N-BRCT repeat of Lig4 interacts with Ku and is required for Lig4 recruitment (Costantini et al. 2007). Consistent with this, deletion of this region abrogated Lig4-X4 colocalization (Figure 3C-D), end synapsis (Figure S3C), and end joining (Figure S3B), even though DNA binding and adenylation activity were retained in this variant (Figure S1B, S3A). Thus, both the Ku and DNA interactions of Lig4 are required for Lig4 colocalization, and neither alone is sufficient.

### A single Lig4 binds both DNA ends at the moment of short-range synapsis

Ligation requires that a single Lig4 binds both DNA ends. We wanted to determine whether such a structure is formed at the moment of short-range synapsis, since this would promote rapid ligation of compatible ends and minimize errors. To this end, we developed a three-color single-molecule FRET assay that simultaneously monitors Lig4 recruitment, Lig4 DNA end binding, and synapsis. As in Figure 2A, we labeled a blunt-ended DNA substrate with Cy3B and Cy5. Additionally, we labeled Lig4 with the near-infrared Cy7 dye (Figure 4A). Thus, using alternating excitation of each dye, this assay monitors Lig4-X4 recruitment (direct Cy7 excitation), short-range synapsis (Cy3B→Cy5 FRET), and Lig4 binding of each DNA end (Cy3B→Cy7 FRET and Cy5→Cy7 FRET) (Figure 4A). Extracts were immunodepleted of endogenous Lig4-X4, supplemented with Cy7-labeled Lig4-X4, and introduced to flow cells containing immobilized Cy3B/Cy5 DNA substrate. Lig4-X4 stoichiometry was estimated by normalizing the Cy7 signal to the average stepwise change in Cy7 signal associated with a binding event (see Methods for details). The example trajectory in Figure 4B shows the following events (orange markers): (1) One Lig4-X4 complex colocalizes (increase in Cy7 signal, bottom panel) and binds the Cy3B DNA end (Cy3B→Cy7 FRET, top panel); (2) a second Lig4-X4 complex colocalizes (increase in Cy7 signal, bottom panel) and binds the Cy5 DNA end (Cy5→Cy7 FRET, middle panel); (3) one of the Lig4-X4 complexes dissociates (decrease in Cy7 signal, bottom panel) and, nearly simultaneously, short-range synapsis occurs (Cy3B→Cy5 FRET, top panel), with the remaining Lig4-X4 complex binding both DNA ends (Cy3B→Cy7 FRET, top panel, and Cy5→Cy7 FRET, middle panel). Consistent with the results presented in Figure S2C, and as shown in this example, we frequently observed Lig4-X4 DNA end binding (Cy3B→Cy7 FRET and/or Cy5→Cy7 FRET) prior to short-range synapsis (Cy3B→Cy5 FRET) (Figure S4A, orange boxes). Across all time points prior to short-range synapsis, DNA molecules exhibited neither end bound by Lig4 (neither Cy3B→Cy7 FRET nor Cy5→Cy7 FRET > 0.25; Figure 4B prior to marker 1 and Figure S4B, red box) in ∼60% of frames; one end bound (either Cy3B→Cy7 FRET or Cy5→Cy7 FRET; Figure 4B, marker 1 and Figure S4B, yellow boxes) in ∼30% of frames; or both ends bound (both Cy3B→Cy7 FRET and Cy5→Cy7 FRET; Figure 4B, marker 2 and Figure S4B, green box) in ∼10% of frames (as in; Figure S4B). Thus, prior to short range synapsis, DNA molecules usually have at most one end bound by Lig4-X4, although occasionally two Lig4-X4 complexes are recruited, with each engaging one end.

**Figure 4:**
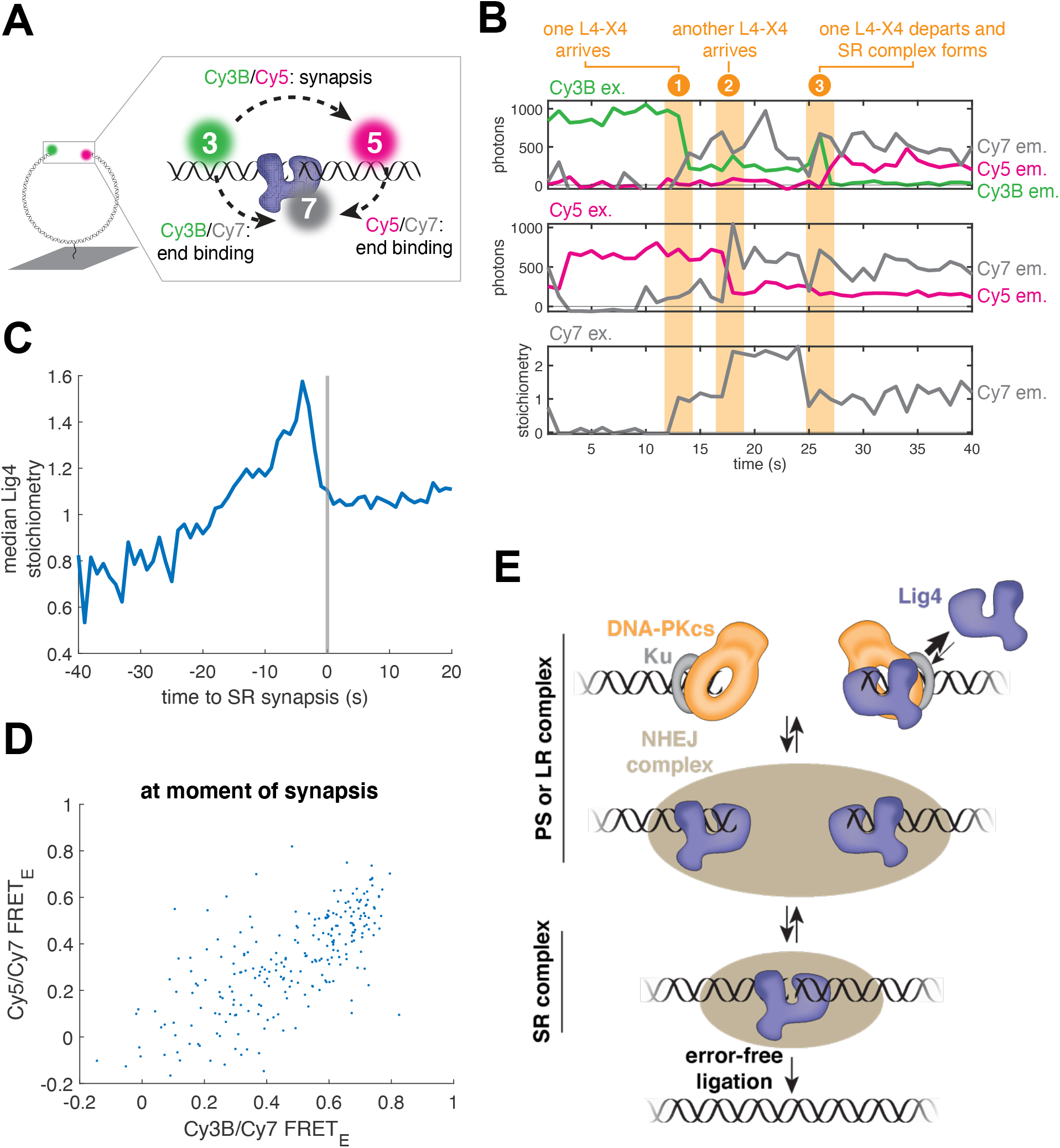
A single Lig4 binds both DNA ends at the moment of short-range synapsis. (A) Cartoon of three-color smFRET assay for synapsis and DNA end binding by Lig4. Green circle: Cy3B fluorphore; magenta circle: Cy5 fluorophore; gray circle: Cy7 fluorophore; dotted arrows: energy transfer. (B) Representative single-molecule trajectories showing Lig4 colocalization, Lig4 end binding, and synapsis. Top panel: Cy3B excitation with Cy3B emission in green, Cy5 emission in magenta, and Cy7 emission in gray; middle panel: Cy5 excitation with Cy5 emission in magenta and Cy7 emission in gray; bottom panel: Cy7 excitation with Cy7 stoichiometry in gray (see Methods for conversion from Cy7 emission to stoichiometry). Orange marker 1: a Lig4 molecule colocalizes (bottom panel) and binds the Cy3B DNA end (Cy3B/Cy7 FRET, top panel); orange marker 2: a second Lig4 molecule colocalizes (bottom panel) and binds the Cy5 DNA end (Cy5/Cy7 FRET, middle panel); orange marker 3: one of the Lig4 molecules dissociates (bottom panel), and the remaining Lig4 molecule binds both DNA ends (Cy3B/Cy7 FRET, top panel; Cy5/Cy7 FRET, middle panel) at the moment of short-range synapsis (Cy3B/Cy5 FRET, top panel). (C) For molecules that underwent short-range synapsis, Cy7 colocalization trajectories (e.g., bottom of panel (B); n = 278 from 8 independent experiments) were aligned, with the onset of short-range synapsis corresponding to t = 0. Blue line shows median Lig4 stoichiometry as a function of time to synapsis. Also see Fig. S4C. (D) Scatter plot depicting Cy3B/Cy7 FRET and Cy5/Cy7 FRET at the moment of shortrange synapsis, as detected by a stepwise increase in Cy3B/Cy5 FRET using the MATLAB ischange function. n = 278 from 8 independent experiments. (E) Model of Lig4-DNA interaction during synapsis. In the presynaptic (PS) complex and/or long-range (LR) synaptic complex, Lig4 transiently binds DNA ends. We postulate that although DNA-PKcs usually occludes DNA ends (top left), DNA ends are transiently accessible and captured by Lig4, which is enriched near DNA ends due to interactions with Ku and potentially other NHEJ factors (see text). Prior to short-range synapsis, both DNA ends are frequently bound by distinct Lig4-X4 complexes, perhaps allowing formation of an XRCC4-XLF-XRCC4 bridge (see text). Immediately before or concurrent with short-range synapsis, one of the Lig4-X4 complexes dissociates such that a single Lig4 binds both DNA ends at the instant of short-range synapsis. Compatible ends can be rapidly joined without errors from this ligation-competent state.

Although recruitment of two L4-X4 complexes was rare overall, we frequently observed two Lig4-X4 complexes 5-10 s before short-range synapsis (as in Figure 4B, marker 2), with one of the two complexes dissociating immediately prior to or concurrent with synapsis (as in Figure 4B, marker 3). To visualize this phenomenon across all observed synapsis events, we aligned trajectories at the instant of short-range synapsis (Figure 4C, gray line) and plotted the median Lig4-X4 stoichiometry (Figure 4C; stoichiometry distributions are shown in Figure S4C). 5-10 s prior to short-range synapsis Lig4-X4 was enriched to a stoichiometry of ∼1.5, before dropping to a stoichiometry of ∼1 at the instant of synapsis. Thus, although the NHEJ complex frequently contains two Lig4-X4 complexes before short-range synapsis, one of these complexes dissociates to leave a single Lig4-X4 at the onset of synapsis. Consistent with this idea, we detected a Lig4-X4 dissociation event within the 10 s period before synapsis for ∼40% of synapsis events (Figure S4D; example in Figure 4B, marker 3), an underestimate due to incomplete Cy7 labeling (∼70%; Figure S4E shows an example trajectory in which no Lig4-X4 dissociation was detected prior to synapsis). Our results are consistent with the transient formation of an XRCC4-XLF-XRCC4 “bridge” (see Discussion) before a single Lig4 mediates short-range synapsis.

We next analyzed the Lig4-DNA interaction at the instant of short-range synapsis, which is marked by an increase in Cy3B→Cy5 FRET (as in Figure 4B, marker 3). As shown in Figure 4B and 4D, most synapsis events were accompanied by high FRET efficiency in both the Cy3B→Cy7 and Cy5→Cy7 channels. Incomplete Cy7 labeling likely accounts for the minor proportion of synapsis events with low Cy3B→Cy7 and Cy5→Cy7 FRET efficiency (example trajectory shown in Figure S4F). Taken together, these results are consistent with a model in which a single Lig4-X4 complex directly binds both DNA ends at the moment of short-range synapsis in a ligation-competent state.

## DISCUSSION

Our findings elucidate the role of Lig4 in the assembly of the short-range synaptic complex, whose formation minimizes mutagenesis by prioritizing ligation over end processing (Figure 4E). Previous work from us and others revealed an essential yet undefined structural role of Lig4 in synapsis (Cottarel et al. 2013; Reid et al. 2015; Graham et al. 2016; Wang et al. 2018; Goff et al. 2022). Here, we demonstrate that Lig4 must directly bind DNA ends to promote short-range synapsis, as mutation of the Lig4 DNA-binding domain blocks short-range synapsis (Figure 2C). Furthermore, three-color smFRET experiments directly visualize that a single Lig4 binds both DNA ends in a ligation-poised state at the instant of short-range synapsis (Figure 4). This moment represents a critical opportunity for error-free ligation of compatible DNA ends, as Lig4 shields DNA ends from processing enzymes that act in the short-range synaptic complex (Stinson et al. 2020). Our results provide a mechanistic basis for NHEJ fidelity, which is observed in multiple model systems (Baumann & West 1998; Labhart 1999; Feldmann et al. 2000; Lin et al. 2013; Waters et al. 2014; Stinson et al. 2020).

We find that the Lig4-DNA interaction is crucial for retention of Lig4-X4 in the NHEJ complex. This interaction complements protein-protein interactions involving direct binding of the N-BRCT and DBD of Lig4 to Ku (Costantini et al. 2007; Chen, Lee, et al. 2021), and the XLF interaction with X4 (Andres et al. 2007). None of these individual interactions is sufficient for Lig4-X4 retention (Figure 3). Instead, Lig4-X4 binding to DNA ends depends on multivalent weak interactions. Such interaction networks allow specific yet reversible binding (Fasting et al. 2012), consistent with the transient Lig4-X4 colocalization we observe. We propose that these Lig4 dynamics are critical for efficient synapsis. Prior to short-range synapsis, both DNA ends are frequently bound by separate Lig4-X4 complexes (Figure 4D). However, one of these Lig4-X4 complexes must dissociate to allow a single Lig4-X4 to bind both DNA ends in the short-range synaptic complex (Figure 4E). Thus, transient Lig4-DNA binding facilitates efficient short-range synapsis.

The short- and long-range synaptic complexes we discovered using single molecule imaging (Graham et al. 2016) have been described at atomic resolution (Chaplin, Hardwick, Liang, et al. 2021; Chaplin, Hardwick, Stavridi, et al. 2021; Chen, Lee, et al. 2021). Here, we reveal additional features of these synaptic states that are not captured in available structures. First, we observe that Lig4 directly binds blunt DNA ends even prior to short-range synapsis (Figures 3,4); in contrast, blunt DNA ends are occluded by DNA-PKcs in X-ray and cryo-EM structures of early-stage NHEJ complexes (Yin et al. 2017; Chaplin, Hardwick, Liang, et al. 2021; Chaplin, Hardwick, Stavridi, et al. 2021; Chen, Lee, et al. 2021; Chen, Xu, et al. 2021; Liu et al. 2022). DNA-PKcs autophosphorylation has been shown to allow access by end-processing enzymes (Ding et al. 2003; Reddy et al. 2004; Goodarzi et al. 2006; Liu et al. 2022), yet we observe robust Lig4-DNA binding even in the presence of DNA-PKcs inhibitor (Figure S2C). We hypothesize that DNA ends are transiently accessible for Lig4 binding prior to DNA-PKcs autophosphorylation. This state may be difficult to resolve by structural methods due to its transient nature. We further propose that the Ku-Lig4^N-BRCT^ interaction, which is strictly required for Lig4-X4 recruitment (Figure 3C,D; (Costantini et al. 2007)), initially enriches Lig4 near DNA ends and allows Lig4 to capture transiently accessible DNA ends. Other Lig4-X4 protein-protein interactions noted above may also play a role in enriching Lig4 near DNA ends. In this way, multiple protein-protein interactions allow Lig4 to access DNA ends prior to short-range synapsis, whereas processing enzymes lacking these interactions are excluded.

In addition, we observe a single Lig4-X4 at the instant of short-range synapsis (Figure 4), whereas a cryo-EM structure of a short-range synaptic complex includes two Lig4-X4 complexes (Chen, Lee, et al. 2021). Consistent with our results, the structure shows one Lig4 binding both DNA ends, whereas the second Lig4 does not appear to bind DNA, and only the BRCT domains are resolved. Our results predict that in the absence of the Lig4-DNA interaction, the second Lig4-X4 would not remain stably associated. It is possible that the presence and/or catalytic activity of DNA-PKcs, which is present in our system but was omitted from the short-range synaptic complex structure, may influence the stability of Lig4-X4. Whereas the cryo-EM study suggests that the presence of two Lig4-X4 complexes is crucial to allow tandem ligation of both DNA strands, our results suggest ligation of the two strands need not be tightly coupled. Consistent with this idea, studies using human cells, human cell extracts, and purified proteins report single-ligation NHEJ intermediates. (Zhou et al. 2010; Pryor et al. 2018; Ma et al. 2004). Our results support a model in which the most critical function of NHEJ is to ligate at least one strand (Waters et al. 2014).

Although we observe a single Lig4-X4 at the instant of short-range synapsis, two Lig4-X4 complexes are frequently present 5-10 s prior to this moment. Notably, XLF is stably recruited to the NHEJ complex within a similar time interval prior to short-range synapsis (Graham et al. 2018). We hypothesize that this Lig4-X4 stoichiometry represents the transient formation of an XRCC4-XLF-XRCC4 “bridge”—a single XLF homodimer interacting with two Lig4-X4 complexes. Uncertainty in the degree of Lig4 labeling precludes us from concluding that two Lig4-X4 complexes are present prior to all or nearly all synapsis events; however, consistent with our single-molecule imaging results, XLF mutations that disrupt one of the two XRCC4 binding sites strongly attenuate short-range synapsis, suggesting that formation of such a bridge is critical for synapsis (Graham et al. 2018). Moreover, this bridge is observed in both long-range and short-range synaptic complex structures (Chen, Lee, et al. 2021; Chaplin, Hardwick, Stavridi, et al. 2021). We speculate that the X4-XLF-X4 bridge may serve multiple functions, including tethering the ends together, stabilizing XLF and/or Lig4-X4 or properly positioning these factors for short-range synapsis.

Our results raise new questions about how NHEJ factor interactions evolve during synapsis and repair. We demonstrate that Lig4 binds DNA ends at the onset of short-range synapsis, and our previous work showed that processing of incompatible ends takes place within the short-range synaptic complex (Stinson et al. 2020). To allow access by processing enzymes, Lig4-X4 likely must release the DNA ends. In the absence of the Lig4-DNA interaction, however, it is unclear how short-range synapsis is maintained. An intriguing possibility is that the XLF homodimer, which is retained following short-range synapsis (Graham et al. 2018) and interacts with Ku on either side of the DSB (Chen, Lee, et al. 2021; Carney et al. 2020), fulfils this role. Further biochemical, single-molecule, and structural studies will be required to fully describe the dynamic intermediates underlying faithful NHEJ.

## Acknowledgments

We thank members of the Loparo and Walter laboratories for helpful discussions and comments on the manuscript. This work was supported by National Institutes of Health grant R01GM115487 (to J.J.L.) and The Howard Hughes Medical Institute (to J.C.W.). J.C.W. is an investigator of the Howard Hughes Medical Institute and an American Cancer Society Research Professor.

## Author contributions

B.M.S. performed all experiments and analyzed the data. S.M.C. designed and performed initial characterization of Lig4 DNA binding mutations. B.M.S, S.M.C., J.C.W., and J.J.L. conceived experiments and wrote the paper.

---

**Figure S1:**
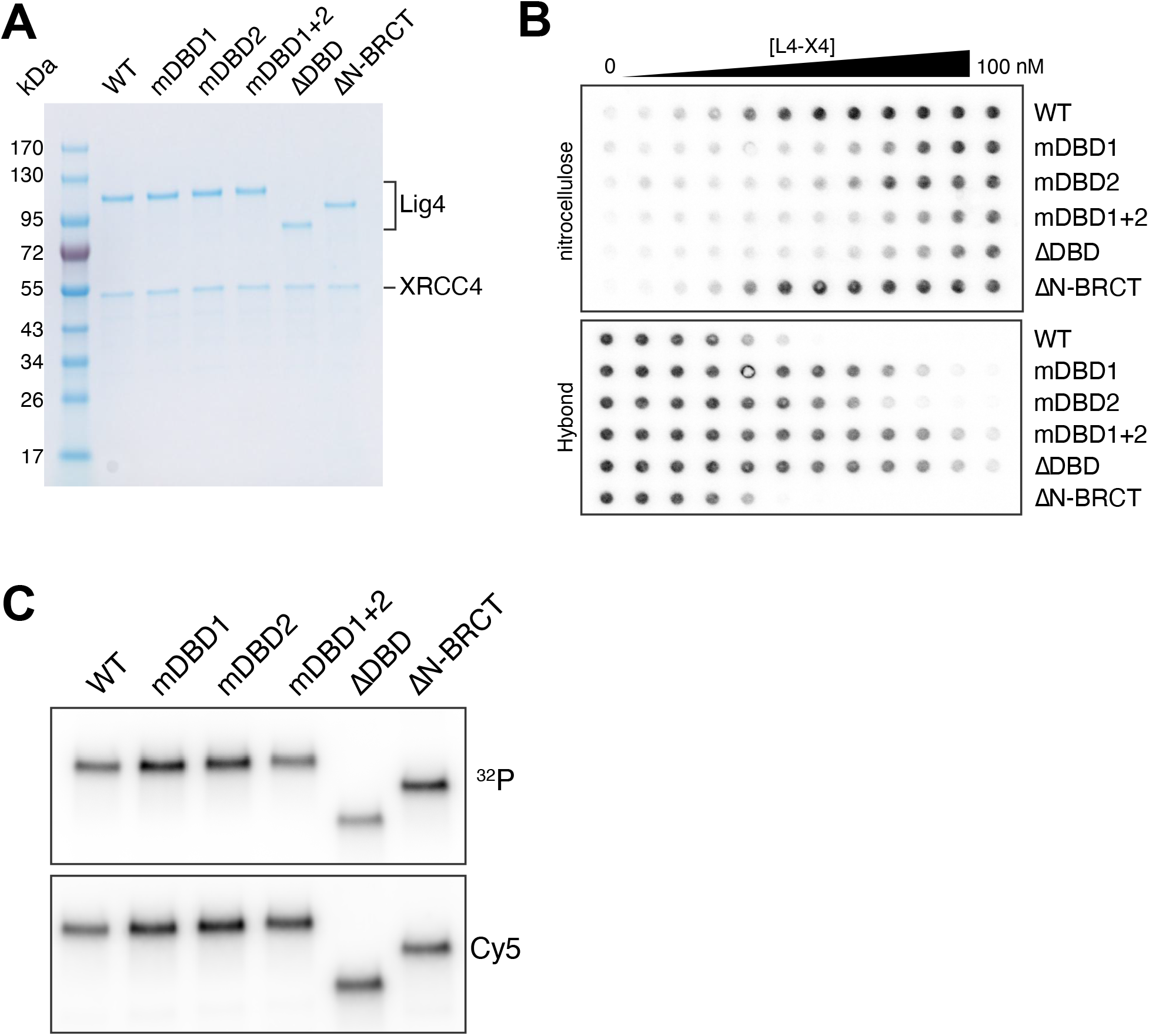
Characterization of Lig4 DNA binding mutants, continued. (A) Coomassie-stained polyacrylamide gel of recombinantly purified Lig4-XRCC4-variants. (B) Representative raw data for filter binding assay shown in Figure 1B and described in Methods. Radiolabeled DNA bound by Lig4 is captured on the nitrocellulose membrane. Free DNA passes through the nitrocellulose membrane and is captured on the Hybond membrane. Lig4-X4 concentration is a two-fold dilution series from 100 nM. (C) Adenylation assay for Lig4 variants, as described in Methods. Cy5-labeled Lig4 variants were de-adenylated with inorganic pyrophosphate and re-adenylated using α-^32^P-ATP and analyzed by PAGE and autoradiography. Top panel shows autoradiogram of ^32^P incorporation; bottom panel shows Cy5 intensity as a loading control. Three independent experiments were performed and representative images are shown.

**Figure S2:**
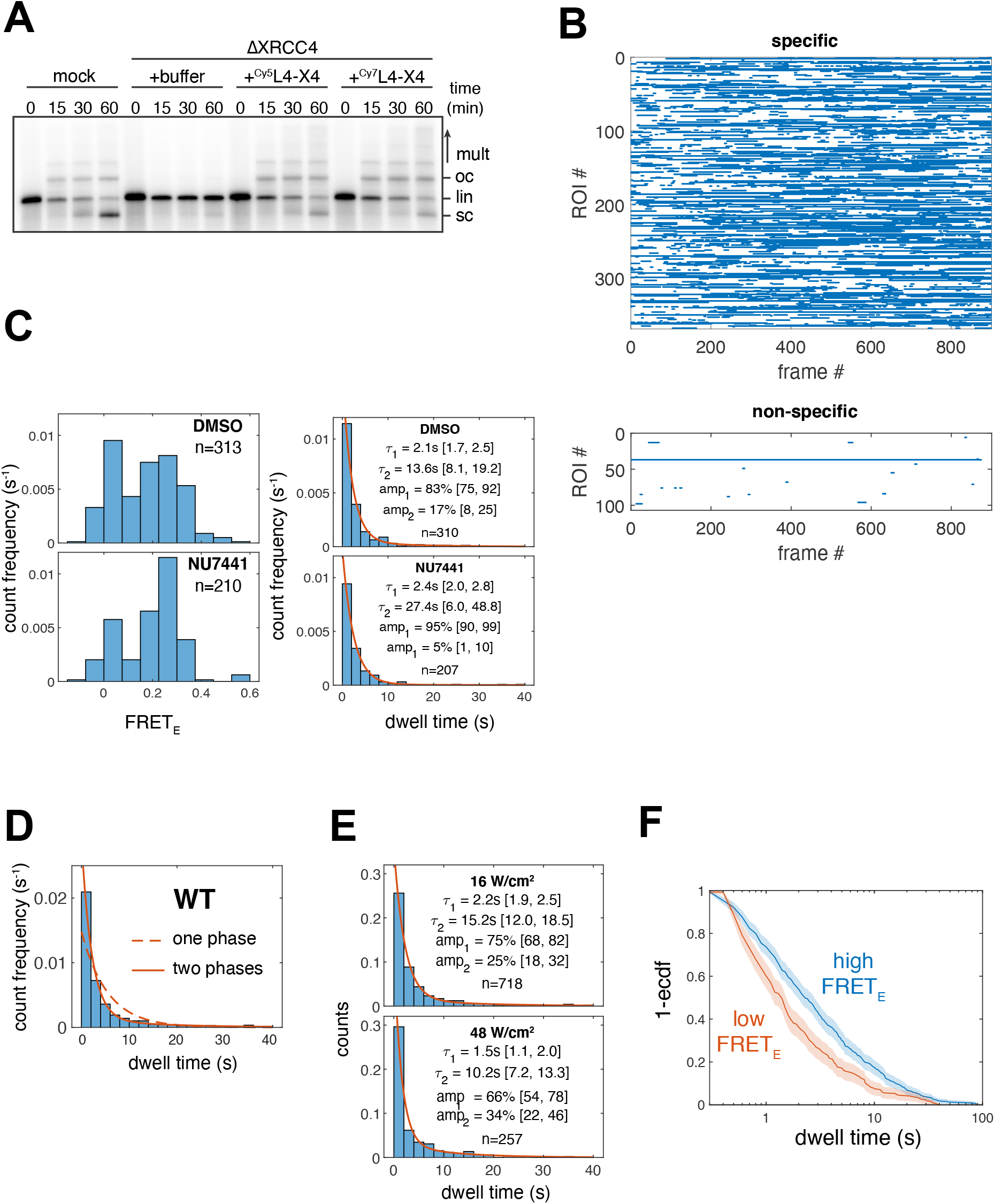
DNA-binding by Lig4 is required for short-range synapsis, continued. (A) Radiolabeled, blunt-ended, linear DNA molecules were added to the indicated extracts, and reaction samples were stopped at the indicated timepoints. Samples were analyzed by agarose gel electrophoresis and autoradiography. lin: linear; sc: supercoiled; oc: open circular; mult: multimers. Three independent experiments were performed and a representative autoradiogram is shown. (B) Rastergrams depicting specific and non-specific ^Cy5^Lig4-X4-^Cy3B^DNA colocalization events from a representative experiment. Each row represents an individual DNA molecule, and blue lines represent times at which Lig4 colocalization was detected. Specific colocalizations occur at regions of interest (ROIs) containing ^Cy3B^DNA signal; non-specific colocalizations occur at ROIs lacking ^Cy3B^DNA signal. (C) WT Lig4 colocalization FRET and dwell time histograms, as in Fig. 3C-E, in extracts treated with DMSO vehicle or 50 μM NU7441 DNA-PKcs inhibitor. Red lines show dwell time distribution fits generated by maximum likelihood estimation in MATLAB using two-term exponential models. Fit parameters: τ, exponential time constant; amp., amplitude. Figures in brackets represent 95% confidence intervals. Data from three independent experiments. (D) WT Lig4 dwell times reproduced from Figure 3D, top panel. Red line shows dwell time distribution fit generated by maximum likelihood estimation in MATLAB using two-term exponential models. Dashed line shows poor fit to a one-term exponential model. (E) Histograms showing dwell time distributions for WT Lig4 at different 641 nm (Cy5 excitation) powers. Upper panel replicates data from top panel of Fig. 3C; lower panel shows histogram at higher 532 nm laser power. Red lines show dwell time distribution fits generated by maximum likelihood estimation in MATLAB using two-term exponential models. Fit parameters: τ, exponential time constant; amp., amplitude. Figures in brackets represent 95% confidence intervals. Data from three independent experiments. (F) WT Lig4 colocalization dwell time partitioned by FRET_E_ (low, <0.1; high, >0.1). ecdf, empirical cumulative distribution function. Data from three independent experiments.

**Figure S3:**
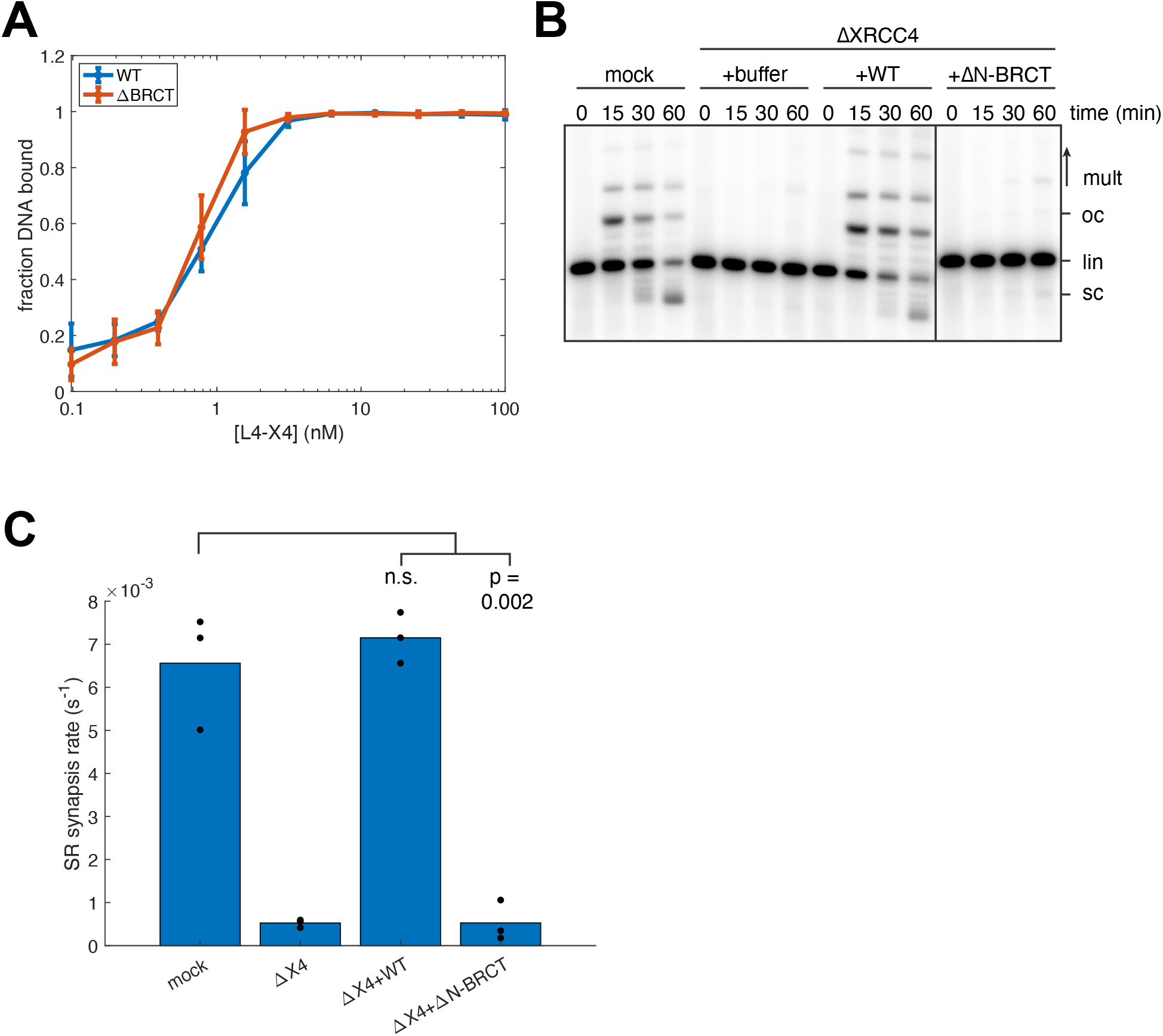
Characterization of Lig4 ΔN-BRCT variant. (A) Filter binding assay for DNA binding by WT and ΔN-BRCT Lig4, as in Fig. 1B. WT data are reproduced from Fig. 1B. (B) End joining assay for WT and ΔN-BRCT Lig4, as in Fig. 1C. Mock, ΔΧ4, and ΔΧ4+WT data are reproduced from Fig. 1C. (C) Short-range synapsis assay for WT and ΔN-BRCT Lig4, as in Fig. 2C. Mock, ΔΧ4, and ΔΧ4+WT data are reproduced from Fig. 2C.

**Figure S4:**
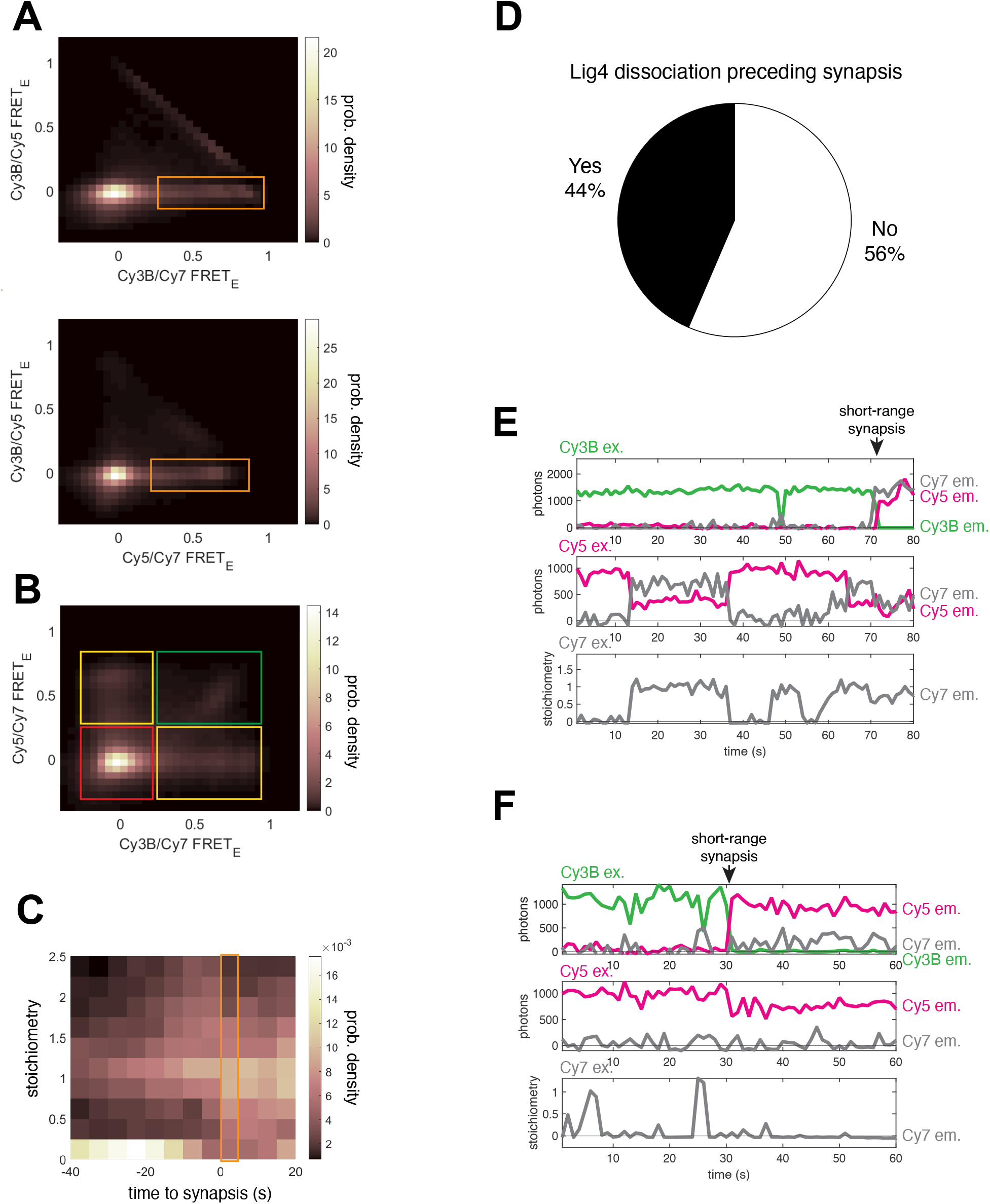
A single Lig4 binds both DNA ends at the moment of short-range synapsis, continued. (A) Heatmaps showing Cy3/Cy5 FRET as a function of Cy3B/Cy7 FRET (top) or Cy5/Cy7 FRET (bottom) for the experiment shown in Figure 4. Orange boxes show Lig4 DNA binding prior to SR synapsis. Heatmaps contain data from all experimental frames in which neither Cy3B nor Cy5 had photobleached. (B) Heatmap showing Cy5/Cy7 FRET as a function of Cy3B/Cy7 FRET for the experiment shown in Figure 4. Heatmap contains data from all experimental frames in which neither Cy3B nor Cy5 had photobleached and Cy3B/Cy5 FRET_E_ was <0.15. Red box: neither Cy3B/Cy7 nor Cy5/Cy7 FRET, indicating neither end is bound by Lig4; yellow boxes: either Cy3B/Cy7 or Cy5/Cy7 FRET, indicating one end is bound by Lig4; green box: both Cy3B/Cy7 and Cy5/Cy7 FRET indicating both ends are bound by a separate Lig4. (C) Heatmap showing stoichiometry distributions relative to onset of synapsis, analogous to Fig. 4D. Orange box highlights stoichiometry distribution at the moment of short-range synapsis. (D) Pie chart depicting the proportion of short-range synapsis events for which a Lig4 dissociation event was detected within a 10 s interval prior to synapsis. Dissociation events were defined as stepwise changes in Cy7 intensity (with 730 nm excitation) of greater than 0.7 stoichiometry units (see Methods). (E) Example trajectory, as in Figure 4B, showing a synapsis event in which dissociation of a second Lig4 molecule was not detected within the 10 s before SR synapsis. (F) Example trajectory, as in Figure 4B, showing a synapsis event in the absence of a labeled Lig4.

## MATERIALS AND METHODS

### Egg extract preparation

High-speed supernatant (HSS) of egg cytosol was prepared as described (Lebofsky et al., 2009).

### Preparation of DNA substrates for end joining

Radiolabeled DNA substrates were prepared as previously described (Graham et al. 2016). In brief, pBMS6, a derivative of pBlueScript, was linearized with Acc65I (New England Biolabs) and resulting 5′ overhangs were filled in using Klenow Fragment polymerase and a mixture of dNTPs including α-^32^P-dATP. The resulting radiolabeled, blunt-ended, linear DNA fragment was recovered using a Qiagen PCR clean-up kit.

Fluorescently labeled DNA substrates were prepared as previously described (Stinson et al. 2020). In brief, pBMS6 was linearized with SphI and AatII (New England Biolabs). The resulting 2977 bp fragment was separated on a 1x TBE agarose gel and recovered by electroelution and ethanol precipitation. Duplex oligonucleotide adapters (Integrated DNA Technologies, Inc.) were ligated to each side of this backbone fragment to generate blunt-ended DNA substrates and purified by agarose gel electrophoresis. (SphI-compatible duplex: CGTACCGC/iAmMC6T/CTAT annealed to ATAGAGCGGTACGCATG; AatII-compatible duplex: CGTACCGC/iAmMC6T/CTAT annealed to CGTGAATACCGCCACGT). DNA was recovered by electroelution and ethanol precipitation and subsequently was treated with Nt.BbvCI (New England Biolabs) to introduce two nicks on the same stand near the middle of the molecule, thereby allowing removal of a 25-mer oligonucleotide. A 10-fold molar excess of an internally biotinylated, 5′-phosphorylated oligonucleotide with the same sequence was then added to the digestion mixture, annealed, and ligated into the gap. DNA substrates were purified by agarose gel electrophoresis and recovered by electroelution and ethanol precipitation. Finally, DNA substrates were phosphorylated with T4 PNK prior to use.

### Preparation of fluorescently-labeled oligonucleotides

Fluorescently-labeled oligonucleotides were prepared as previously described (Stinson et al. 2020). In brief, amino-modified synthetic oligonucleotides were reacted with NHS-ester fluorophore derivatives, purified by denaturing PAGE, and recovered using a crush-and-soak procedure followed by ethanol precipitation.

### Protein expression and purification

The following protocol was used for expression and purification of all L4-X4 variants. Both sets of L4-X4 variants (unlabeled and Cy5-labeled) were expressed and purified in parallel. pETDuet-1 plasmids were constructed encoding Xenopus Lig4 and XRCC4. Lig4 contained a TEV-cleavable N-terminal His_6_ tag and a 3C protease cleavable C-terminal TwinStrep tag. XRCC4 was untagged. Expression plasmids were transformed into BL21 cells and plated on LB-agar plates containing 100 μg/mL ampicillin. Single colonies were used to inoculate 5 mL LB cultures supplemented with 100 μg/mL ampicillin, which were grown overnight at 37 °C. These starter cultures were added to 250 mL Terrific Broth supplemented with 100 μg/mL ampicillin, and cultures were shaken at 37 °C to an optical density of 1.2. Cultures were moved to a shaker at 16 °C, and 1 mM IPTG was added to induce expression overnight. Cultures were harvested by centrifugation and resuspended in ice-cold 20 mL lysis/wash buffer (20 mM Tris, pH 8,0; 400 mM NaCl; 10 mM imidazole; 1 mM DTT; 10% glycerol). A Roche cOmplete Protease Inhibitor Cocktail (EDTA free) tablet was added to the resuspension, and cells were lysed by sonication on ice. The lysate was centrifuged at 50,000g for 60 min at 4 °C, and the supernatant was added to 0.5 mL (bed volume) Ni-NTA resin (Qiagen) equilibrated in wash/lysis buffer. The resulting mixture was rotated at 4 °C for 60 min and then poured into a 1 mL polypropylene column (Qiagen). The resin was washed three times with 5 mL lysis/wash buffer and eluted in 0.25 mL fractions with Ni-NTA elution buffer (composition same as lysis/wash buffer, but with 250 mM imidazole). Fractions were analyzed by SDS-PAGE, and fractions containing Lig4-XRCC4 were applied to a 0.5 mL (bed volume) Streptactin XT column (IBA Biosciences) equilibrated in lysis/wash buffer. The Streptactin XT column was washed three times with lysis/wash buffer and eluted in 0.5 mL fractions with 1x BXT buffer (IBA Biosciences). Fractions containing Lig4-XRCC4 were concentrated with a 10 kDa MWCO Amicon centrifugal filter unit and loaded onto a Superdex200 Increase 10/300 GL (Cytiva) column equilibrated in SEC buffer (20 mM HEPES, pH 7.5; 150 mM NaCl; 1 mM DTT; 10% glycerol). Fractions containing pure Lig4-XRCC4 were concentrated with a 10 kDa MWCO Amicon centrifugal filter unit, aliquoted, flash frozen in liquid nitrogen, and stored at -80 °C. Protein concentrations were estimated by absorbance at 280 nm.

For fluorescently-labeled Lig4, the 11 amino acid ybbR tag (Yin et al. 2006) was inserted between the TEV recognition site and Lig4 CDS, and labeling took place prior to gel filtration chromatography. Concentrated fractions from the Streptactin column were supplemented with final concentrations of 10% glycerol (v/v), 10 mM MgCl2, 5 μM Sfp synthase (plasmid obtained from Addgene (pET-Sfp, #159617) and purified as described (Yin et al. 2006)), and 100 μM CoA-Cy5 or CoA-Cy7 (prepared as described below). Labeling reactions were carried out in the dark overnight at 4 °C, and labeling reactions were loaded directly onto a Superdex200 Increase column as described above. Labeling efficiencies were 60-80% as estimated by absorbance at the dye absorbance maximum.

### Preparation of Coenzyme A-fluorophore conjugates

CoA-dye conjugates were prepared essentially as described (Yin et al. 2006), with a modified purification procedure. Briefly, 0.5 mg maleimide-functionalized sulfonated Cy5 or Cy7 (Lumiprobe cat. # 13320 or 15320) in 125 μL DMSO was added to 1 mg CoA trilithium salt in 375 μL 100 mM HEPES, pH 7.0. The reaction was allowed to proceed in the dark at room temperature for 1 hour. The reaction mixture was loaded onto two 1 mL HiTrapQ columns (Cytiva) connected in series and equilibrated in 10% acetonitrile (v/v). CoA conjugates were purified using a 0-20% gradient over 20 column volumes (A: 10% acetonitrile; B: 3 M LiCl), with desired conjugates eluting as the final peak of the chromatogram. Fractions from this peak was pooled and CoA conjugates were precipitated with 20 volumes cold acetone and collected by centrifugation. Pellets were washed with cold acetone and resuspended in 10 mM Tris, pH 8.0 CoA conjugate concentrations were estimated by absorbance at the dye absorbance maximum.

### Filter binding assay for DNA binding

Filter binding experiments used a radiolabeled 1 kb circular DNA substrate to preclude complications from ligation of the DNA substrate. As described above in “Preparation of DNA substrates for end joining,” pBMS6 was linearized with Acc65I (New England Biolabs) and resulting 5′ overhangs were filled in using Klenow Fragment polymerase and a mixture of dNTPs including α-^32^P-dATP. The resulting radiolabeled, blunt-ended, linear DNA fragment was recovered using a Qiagen PCR clean-up kit, diluted to ∼1 nM in 1x T4 DNA ligase reaction buffer (New England Biolabs), and treated with T4 DNA ligase to circularize the DNA. Ligation reactions were concentrated using a 10 kDa MWCO Amicon centrifugal filter unit and separated on a 1% agarose gel containing ethidium bromide by electrophoresis. The supercoiled 1 kb band was excised and purified using a Qiagen gel extraction kit.

The filter binding assay protocol was modified from (Wong & Lohman 1993). Hybond-N+ (Cytiva) and nitrocellulose (Whatman) membranes were equilibrated in a 1:3 mixture of SEC buffer:FB buffer (50 mM Tris, pH 8.0; 1 mM MgCl_2_). Membranes were assembled in a Biorad Bio-Dot apparatus with the nitrocellulose membrane on top of the Hybond membrane. Serial 2-fold dilutions of Lig4-X4 variants were prepared in SEC buffer in a multi-well plate, and the DNA substrate was diluted to 6 pg/mL in FB buffer. Binding reactions were initiated by adding 15 μL of the DNA mixture to 5 μL of the protein mixture and were allowed to proceed for 5 min at room temperature. Reactions were transferred to the Bio-Dot apparatus with a multi-channel pipette, and vacuum was immediately applied to draw samples through the membranes. Each well was washed twice by adding 100 μL ice-cold 1:3 SEC buffer:FB buffer and rapidly applying vacuum. Membranes were dried on a vacuum gel drier and exposed to storage phosphor screens for autoradiography and imaged with a Typhoon FLA 7000 imager (GE Healthcare).

### Lig4 autoadenylation assay

Autoadenylation assays were performed as described previously (Graham et al. 2016) with slight modifications. In a 10 μL reaction volume, 50 nM Lig4-X4 variants were treated with 50 μM inorganic pyrophosphate in adenylation buffer (60 mM Tris, pH 8.0; 10 mM MgCl_2_; 5 mM DTT; 5 μg/mL BSA; 10% glycerol) for 2 min at room temperature to de-adenylate Lig4. α-^32^P-ATP (3000Ci/mmol, 10mCi/ml; Perkin Elmer) was diluted 4-fold in adenylation buffer, and 1 μL was added to each sample. Adenylation was allowed to proceed for 10 min at room temperature before being quenched with one volume of 2x Laemmli sample buffer. Samples were separated by SDS-PAGE, and gels were fixed with 10% methanol/10% acetic acid before being dried on a vacuum gel drier, exposed to storage phosphor screens for autoradiography, and imaged with a Typhoon FLA 7000 imager (GE Healthcare).

### Antibodies and immunodepletion

The rabbit polyclonal antibody raised against XRCC4 was previously described (Graham et al. 2016). XRCC4 immunodepletions were carried out using the following protocol: 3 volumes of 1 mg/mL affinity-purified antibody was gently rotated with 1 volume Protein A Sepharose beads (GE Healthcare) overnight at 4°C or 1 hour at room temperature. Beads were washed extensively with ELBS (2.5 mM MgCl2, 50 mM KCl, 10 mM HEPES, pH 7.7, 0.25 M sucrose), and ten volumes of egg extract containing 7.5 ng/μL nocodazole were immunodepleted by two rounds of gentle rotation with one volume of antibody-bound beads for 20 min at room temperature. Immunodepleted extracts were either used immediately or aliquoted and flash-frozen in liquid nitrogen.

### Ensemble NHEJ assays

Ensemble NHEJ assays were conducted at room temperature. Egg extracts were supplemented with the following (final concentration indicated in parentheses): pBMS6 (30 ng/μL); ATP (3 mM); phosphocreatine (15 mM); and creatine phosphokinase (0.01 mg/mL; Sigma). Joining reactions were initiated by addition of 2.5 ng/μL radiolabeled linear DNA substrate (final concentration, prepared as described above).

For analysis by agarose gel electrophoresis, samples were withdrawn at the indicated times and mixed with a 2.5 volumes agarose stop solution (80 mM Tris, pH 8.0, 8 mM EDTA, 0.13% phosphoric acid, 10% Ficoll, 5% SDS, 0.2% bromophenol blue). Samples were treated with Proteinase K (1.4 mg/mL final concentration) for 60 min at 37°C or room temperature overnight, and products were separated by electrophoresis on a 1x TBE 0.8% agarose gel. Gels were dried under vacuum on a HyBond N+ membrane (GE Healthcare) and exposed to a storage phosphor screen, which was imaged with a Typhoon FLA 7000 imager (GE Healthcare).

### Single-molecule microscope, chamber preparation, and general protocol

Samples were imaged with a through-objective TIRF microscope built around an Olympus IX-71 inverted microscope body (Graham et al. 2017). 532 nm, 641 nm, and 730 nm laser beams (Coherent Sapphire 532, Cube 641, and OBIS 730, respectively) expanded, combined with dichroic mirrors, expanded again, and focused on the rear focal plane of an oil-immersion objective (Olympus UPlanSApo, 100×; NA, 1.40). The 730 nm laser beam was passed through a Chroma ZET730/10x clean-up filter prior to expansion. Lasers were switched on and off using Uniblitz V14 shutters. The focusing lens was placed on a vertical translation stage to permit manual adjustment of the TIRF angle. Emission light was separated from excitation light with a multipass dichroic mirror (ZT405/488/532/640/730rpc-uf2and; Chroma) mounted in an Olympus BX filter cube, and laser lines were further attenuated with a ZET405/488/532/640m emission filter (Chroma) and StopLine 488/532/635 (Semrock) and ZET730nf (Chroma) notch filters. A home-built beamsplitter (Graham et al. 2017) was used to separate Cy3B emission from Cy5/Cy7 emission using a Chroma T640lpxr dichroic. Chroma ET650sp and 488/532m emission filters were placed in the Cy3B emission path, and a filter wheel was placed in the Cy5/7 emission path to select for Cy5 emission (ET700/75m; Chroma) or Cy7 emission (ET811/80m; Chroma). These two channels were imaged on separate halves of an electron-multiplying charge-coupled device camera (Hamamatsu, ImageEM 9100-13), which was operated at maximum EM gain. A motorized microstage (Mad City Labs) was used to position the sample and move between fields of view.

Microfluidic chambers were constructed in the following manner: a Dremel tool with a diamond-tipped rotary bit was used to drill two holes 10 mm apart in a glass microscope slide; PE20 tubing was inserted into one hole and PE60 tubing into the other (Intramedic), and the tubing was cut flush on one side of the slide and fixed in place with epoxy (Devcon) on the other; double-sided SecureSeal Adhesive Sheet (Grace Bio-Labs), into which a 1.5 × 12 mm channel had been cut, was placed on the non-tubing side of the slide, aligning the channel with the holes in the slide. A glass coverslip, functionalized with a mixture of methoxypolyethylene glycol-succinimidyl valerate, MW 5,000 (mPEG-SVA-5000; Laysan Bio, Inc.) and biotin-methoxypolyethylene glycol-succinimidyl valerate, MW 5,000 (biotin-PEG-SVA-5000; Laysan Bio, Inc.) as previously described (Graham et al., 2017), was then placed on the second side of the adhesive sheet, and the edges of the coverslip were sealed with epoxy.

Single-molecule experiments were generally performed as follows, with modifications noted in the following sections. Solutions were drawn into the chamber by attaching a 1 mL syringe to the PE60 tubing. Flow cells were incubated with 1 mg/mL streptavidin (Sigma) in PBS for ∼2 min. Unbound streptavidin was washed out with ELBS and biotinylated DNA substrates were incubated in the channel at a concentration yielding appropriate surface density (typically ∼ 1 nM, diluted in ELBS). Unbound DNA was washed out with ELBS and experiments were performed as indicated below. Extracts were supplemented with pBMS6 (100 ng/μL); ATP (3 mM); phosphocreatine (15 mM); creatine phosphokinase (0.01 mg/mL; Sigma); protocatechuic acid (PCA; 5 mM); protocatechuate-3,4-dioxygenase (PCD; 0.1 μM); ascorbic acid (1 mM); and methyl viologen (1 mM). PCA/PCD constitute an oxygen scavenging system (Aitken et al. 2008) and ascorbic acid/methyl viologen suppress fluorophore blinking (Stein et al. 2012).

### Two-color single-molecule assay for short-range synapsis

The two-color single-molecule assay for short-range synapsis (Fig. 2 and S3C) was performed as previously described with slight modifications (Graham et al. 2016). The biotinylated, Cy3B/Cy5 DNA substrate was immobilized on a glass coverslip in a microfluidic chamber as described above. Extracts were immunodepleted of XRCC4 and supplemented with SEC buffer or 50 nM unlabeled Lig4-X4 variant, as well as pBMS6 (100 ng/μL); ATP (3 mM); phosphocreatine (15 mM); creatine phosphokinase (0.01 mg/mL; Sigma); protocatechuic acid (PCA; 5 mM); protocatechuate-3,4-dioxygenase (PCD; 0.1 μM); ascorbic acid (1 mM); and methyl viologen (1 mM). Extract was introduced to the chamber and images were taken continuously at a rate of 2 frame/s, alternating between one frame of 532 nm excitation and one frame of 641 nm excitation. Each field-of-view (FOV) was imaged for 2 min, and five FOVs were imaged per experiment. Surface laser power density was measured through the objective with epi-illumination using a Coherent FieldMate power meter with an OP-2 VIS detector (532 nm: 8 W/cm^2^; 641 nm: 4 W/cm^2^). Synapsis events were automatically detected as stepwise increases in FRET_E_ using a custom MATLAB script. Synapsis rates were calculated as the number of synapsis events divided by the total time DNA molecules were unsynapsed with both fluorophores intact.

### Two-color single-molecule assay for Lig4 colocalization and DNA binding

The two-color single-molecule assay for Lig4 colocalization and DNA binding (Fig. 3, S2) was performed as described in the preceding section with the following modifications: DNA substrate was labeled with Cy3B on both ends; extracts were supplemented with 10 nM Cy5-labeled Lig4-X4 variants; images were taken continuously at a rate of 20 frame/s, alternating between one frame of 532 nm excitation and one frame of 641 nm excitation; one FOV was imaged for 6 min per experiment; laser powers were 16 W/cm^2^ (532 nm) and 16 W/cm^2^ (641 nm). For the experiment in Fig. S3C, 641 nm laser power was increased to 48 mW/cm^2^. To ensure analyzed DNA spots had two intact Cy3B fluorophores, minimum DNA spot intensity was set at a threshold determined by the intensities of spots showing two Cy3B photobleaching events. See “Single-molecule data analysis” below for further details of colocalization and FRET analysis.

### Three-color single-molecule assay for synapsis, Lig4 colocalization/DNA binding

The three-color single-molecule assay for synapsis and Lig4 colocalization/DNA binding (Fig. 4, S4) was performed as in the two-color single-molecule assay for short-range synapsis with the following modifications: extracts were supplemented with 20 nM Cy7-labeled wild-type Lig4-X4; images were taken continuously at a rate of 5 frame/s with the following iterated sequence: 532 nm ex. with Cy3B/Cy5 emission, 532 nm ex. with Cy3B/Cy7 emission, 641 nm ex. with Cy5 emission, 641 nm ex. with Cy7 emission, 730 nm ex. with Cy7 emission; laser powers were 40 W/cm^2^ (532 nm), 40 W/cm^2^ (641 nm), and 300 W/cm^2^ (730 nm).

To convert Cy7-Lig4 intensities to stoichiometries, stepwise changes (corresponding to Lig4 binding/unbinding events) were detected in the Cy7 ex./Cy7 em. trajectory for each DNA molecule using the MATLAB `ischange` function. Cy7 intensities were normalized by the average step intensity for each molecule to generate a Lig4 stoichiometry estimate. Synapsis events were automatically detected as stepwise increases in Cy3B/Cy5 FRET_E_ using a custom MATLAB script.

### Single-molecule data analysis

A publicly available automated analysis pipeline (Smith et al. 2019) was used for spot detection and determination of local background-corrected fluorescence intensities and Lig4 colocalization events. To determine Lig4 colocalization events within the pipeline, thresholds were set for minimum Lig4 spot intensity and displacement from the DNA spot, and events were required to last for at least two frames with at least two frames between successive colocalization events. Thresholds were chosen to minimize non-specific colocalization events at ROIs lacking DNA signal. Fluorescence intensities were exported from the pipeline and corrected for bleed through from donor channel to acceptor channel, direct excitation of acceptor fluorophore by donor excitation laser, and differences in fluorophore quantum yield and detection efficiency (gamma factor; (Roy et al. 2008)). Corrected intensities were used to calculate apparent FRET efficiencies. For two-color experiments, Cy3B/Cy5 FRET_E_ was calculated as Cy5 intensity upon Cy3B excitation divided by the sum of Cy3B and Cy5 intensity upon Cy3B excitation (E_35_ = I_35_/(I_3_+I_35_)). For three-color experiments, apparent FRET efficiencies were calculated as follows: E_35_ = I_35_/(I_3_+I_35_+ I_37_); E_37_ = I_37_/(I_3_+I_35_+ I_37_); E_57_ = I_57_/(I_5_+I_57_).

